# Myosin-9b Controls Epithelial Brush Border Architecture through Motility-Dependent RhoA Signaling

**DOI:** 10.64898/2026.09.08.750145

**Authors:** Emma C. Murray, Namya Manoj, Lynn Ziegler, Gillian M. Hodge, Shane Fraher, Leighton S. Lee, Lucas Code, Steven A. Lewis, Christine E. Schaner Tooley, Yongho Bae, John Konen, Jonathan E. Bard, Andrew T. Lombardo

## Abstract

Genetic variations in the *MYO9B* gene have been associated with Crohn’s disease, celiac disease, and ulcerative colitis. These diseases have been characterized as primarily immune disorders. However, the overall molecular basis for the influence of Myo9b in these diseases remains poorly understood. Using *in vivo* small intestine ileum and human cell culture models, we identify a molecular function for Myo9b in the regulation of epithelial brush border microvilli. Using live-cell super-resolution microscopy, we characterize the motility of Myo9b as it moves toward enriched puncta at the tips of microvilli and visualize its direct regulation of small GTPase signaling using an active RhoA biosensor. In Myo9b knockout cells, microvilli abundance and dynamics are altered, but the cells ultimately maintain the presence of microvilli and the appropriate incorporation of microvilli specific cytoskeletal to membrane regulators such as Ezrin. Alternatively, expression of the Myo9b-S1011A disease variant as the only genetic copy in human cells results in a total loss of microvilli and Ezrin apical localization. These results indicate that Myo9b is a critical regulator of epithelial cell morphology and microvilli. Further, our data establish that the S1011A disease variant disrupts microvilli in human cells, suggesting a potential mechanistic link to its involvement in disease states.

**Significance Statement:** Inflammatory bowel diseases (IBD) are characterized by intestinal barrier dysfunction. However, it is unknown whether the barrier dysfunction is a primary cause of the disease or a result of the disease-induced immune response. Here, we show that the protein Myosin 9b is required to regulate the epithelial brush border microvilli in human cell culture models, where its motile properties localize its signaling domain to microvilli tips. Introducing Myo9b disease variants into cells results in the loss of proper epithelial cell morphology. Our data suggest a potential mechanism where disease causing Myo9b mutants produce a functionally disruptive protein, contributing to the destruction of the intestinal barrier in disease states.

## Introduction

Inflammatory bowel diseases (IBD) are characterized by both an autoimmune response and dysfunction of the epithelial barrier, which allows improper passage of antigens, bacteria, and molecules across the barrier, resulting in an inappropriate immune response (1). Myosin 9b (Myo9b) provides a potential link to IBD, as single nucleotide polymorphisms (SNPs) in the *MYO9B* gene have been identified through multiple genetic screens to IBD (2-6).

Myo9b is an actin-based molecular motor characterized to be one of only a handful of motorized signaling molecules within the human genome (7, 8). The C-terminal tail domain of Myo9b encodes a GTPase activating protein (GAP) domain (9). Myo9b’s GAP activity is specific for the negative regulation of the small signaling GTPase, RhoA, by accelerating the transition between active GTP-bound toward the inactive GDP-bound conformation (10, 11). Single molecule biophysical studies have characterized the N-terminal myosin motor domain of Myo9b as a single-headed processive motor that moves towards the plus end of actin filaments (12, 13). The myosin motor domain utilizes the hydrolysis of ATP to produce force and motion on actin filaments. Unlike nearly all other myosins, Myo9b possesses an additional ATP independent actin binding motif, allowing it to remain tethered to actin separate from its ATPase activity (9, 14, 15). The single molecule motility of Myo9b has been characterized *in vitro* and is measured to move at 19.6 ± 7 nm/s (12). However, the motility of Myo9b has never been characterized in a cellular context, and its slow motility *in vitro* has left the function of the motor’s activity *in vivo* uncertain. Therefore, Myo9b possesses at least two separate means of regulating the actin cytoskeleton, through direct interaction via its motor domain and through inactivation of RhoA through its GAP domain (16).

Myo9b is highly expressed in immune cells, with additional expression found in epithelial cells (17, 18). In macrophages, leukocytes, and dendritic cells, reduction of Myo9b results in morphological defects and reduced cellular motility (19-23). Collectively, prior studies have identified that Myo9b functions at sites of active actin polymerization in cell projections, such as extending lamellipodia and filopodial tips, where it regulates RhoA locally (8, 21, 24). In epithelial cells, knockdown of Myo9b impairs wound healing and tight junction formation (23). However, little is known about the specific cytoskeletal structures that Myo9b regulates in epithelial cells or why genetic variants in Myo9b exacerbate IBD.

Epithelial cells of the gut, kidney, and placenta express a continuous layer of microvilli, finger-like projections that increase surface area for absorption and serve as part of the barrier defense at the apical, lumen-facing membrane. In IBD states, microvilli are severely disrupted, where the core actin cytoskeletal bundles that support microvilli are lost or severely depleted (25). In this study, we identify several novel functions for Myo9b in the regulation of apical epithelial morphology: Myo9b localizes the tips of mouse ileum brush boarder microvilli *in vivo*. Super-resolution live imaging in human epithelial cells resolves Myo9b motility along actin bundles of microvilli at velocities exceeding that of the single *in vitro* motor and accumulating at the tips. Myo9b requires both a functional motor domain and RhoGAP domain for proper regulation of microvilli and epithelial cell morphology. Finally, expression of the S1011A disease variant (4-6) as the only copy in human cells results in mislocalization of the variant Myo9b to junctions, resulting in near total loss of microvilli. Collectively, these results represent the first step toward characterizing the specific role of Myo9b in barrier defects of IBD in human cells.

## Results

### Myo9b is Expressed in Epithelial Cells and is Enriched at the Apical Surface

To determine the tissue specificity and expression level of Myo9b in epithelial cells, Western blot analysis was performed on placenta, kidney, and colorectal cultured epithelial cell lysates. The Western blots showed that Jeg-3 placental, Caco-2 colorectal, DLD-1 colorectal, and epithelial-like HeLa cervical cells all expressed Myo9b. All tested lysates had two bands of Myo9b, (Fig 1A, B) arising from the two isoforms produced by alternative splicing. (26). We quantified the total expression of Myo9b and determined that HeLa cells and Jeg-3 cells had the highest relative levels of Myo9b expression (Fig. 1A, B). However, Myo9b expression was not detectable in kidney PK-1 cells (Fig. 1A, B), indicating that Myo9b is broadly expressed in multiple but not all epithelial cells.

**Figure 1.**
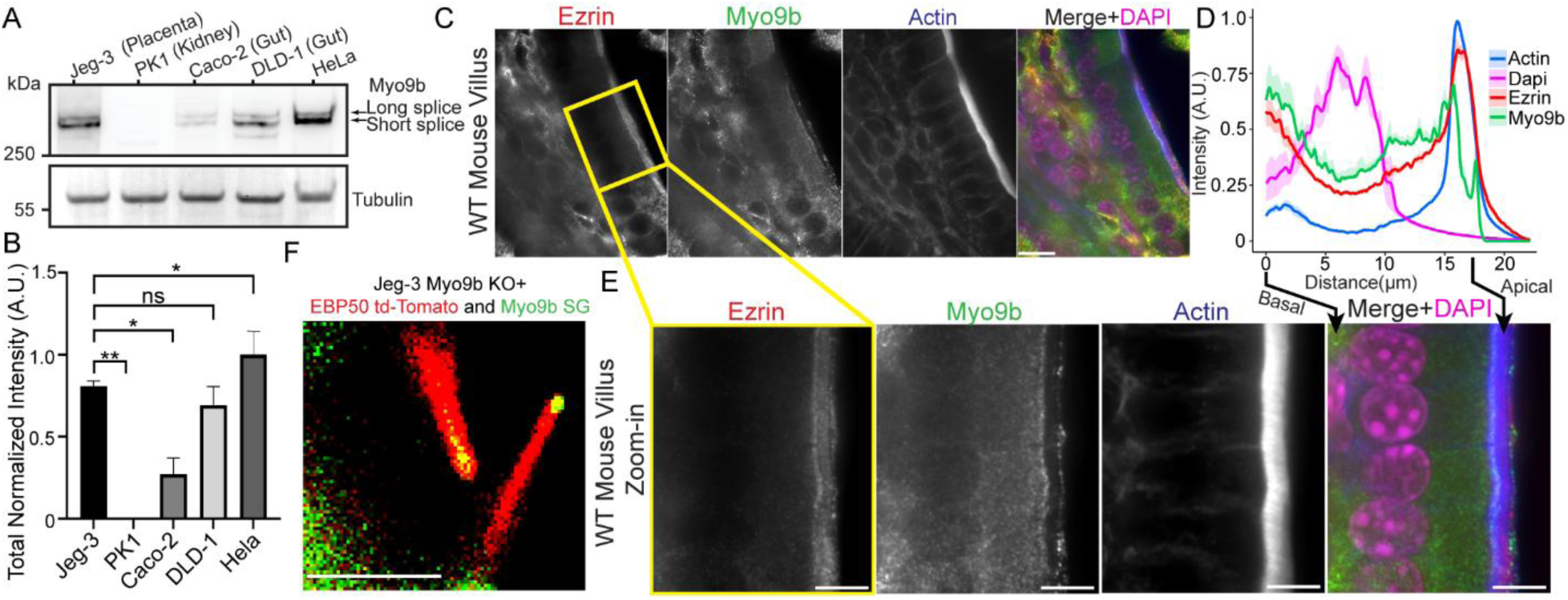
Characterization of Myo9b in Epithelial Cells and WT Mouse Intestinal Tissue. A) Western blot of Myo9b and tubulin in WT Jeg-3, WT PK-1, WT Caco-2, WT DLD-1, and WT HeLa cells. Long splice and short splice indicate the two Myo9b isoforms. B) Quantification of the normalized intensity of Myo9b signal the western blot from (A). Bars indicate Mean ± SEM; Significance by unpaired t-test (n=3, *=p ≤ 0.05, **=p ≤ 0.01, ns = not significant). C) Spinning disk confocal SORA immunofluorescent imaging of WT mouse intestinal villi tissue stained for anti-ezrin, anti-Myo9b, phalloidin (actin), and DAPI. Scale bar 10 μm. D) Fluorescence intensity profiles of actin, DAPI, ezrin, and Myo9b. Intensity profiles extend from the basal membrane through the nucleus to the intestinal lumen for each measured cell. The line for each intensity profile indicates the sliding-average intensity across measured cells, and the shaded region around the line represents the sliding standard error of the average. The colocalized ezrin and actin peak represents the average location of the microvilli, and the DAPI peak is the average location of the nucleus. E) Zoom in of boxed area in (C). Scale bar 5 μm. F) First frame of Video 1 at t=0 showing individual microvilli co-stained for tdTomato-EBP50 (Red) and StayGold-Myo9b (green). Scale bar 2 μm.

We hypothesized that Myo9b would be primarily localized to regions of dynamic, polymerizing actin cytoskeleton within the cell, as had been observed in melanoma cells, osteocytes, motile epithelial cells, and macrophages. (8, 21, 23, 24). However, the specific subcellular domains of localization in non-motile human epithelial cells forming the columnar monolayered brush border of the gut and placenta were unknown. Super Resolution by Optical Pixel Reassignment (SORA) confocal imaging of WT mouse ileum showed Myo9b staining diffusely distributed throughout the cytoplasm with enrichment at the basal plasma membrane of cells, as previously reported (8, 10) (Fig. 1C-E). Interestingly, at the apical surface, Myo9b localized to two enriched regions at the base and tip of the brush border microvilli but was reduced throughout the ezrin-stained microvilli core (Fig. 1C-E). Quantification of the SORA fluorescent intensity profiles revealed that the Myo9b signal straddles both the ezrin and actin fluorescent intensity peaks of the microvilli core bundles of the brush border, indicating Myo9b localizes to the base of the microvilli (terminal web) and to the tips of the microvilli (Fig. 1D). To validate this finding in our cell culture model system, we transiently transfected Jeg-3 cells with a fluorescently tagged StayGold-Myo9b (SG-Myo9b) and the microvilli marker tdTomato-EBP50 (27). Using two-color live SORA confocal microscopy, we identified that SG-Myo9b localized to the tips of microvilli in Jeg-3 cells (Fig. 1F, Movie 1). These data indicate that in the epithelial cells, both *in vivo* mouse ileum and cultured human epithelial Jeg-3 cells, Myo9b localizes to the terminal web and tips of microvilli absent from the microvilli core.

### CRISPR Knockout of Myo9b in Human Epithelial Cells Results in Morphological and Actin Cytoskeletal Defects

To investigate the effect of the loss of Myo9b on cell morphology and cytoskeletal regulation, CRISPR/Cas9 was used to genetically knock out (KO) endogenous Myo9b within Jeg-3 cells. We validated the Myo9b KO by isolating multiple clones of cells lacking Myo9b expression as measured by western blot analysis (Fig. 2A). The validation of a Myo9b KO cell line was attempted in multiple epithelial cell lines of differing tissue types including HeLa cells, as they contain similar levels of Myo9b expression compared to Jeg-3 cells (Fig. 1A). However, during the generation of the Myo9b KO in HeLa cells, the cells lost their ability to execute successful cell division resulting in single giant multi-nucleated syncytial cells making culturing and validation impossible (Fig. S1).

**Figure 2.**
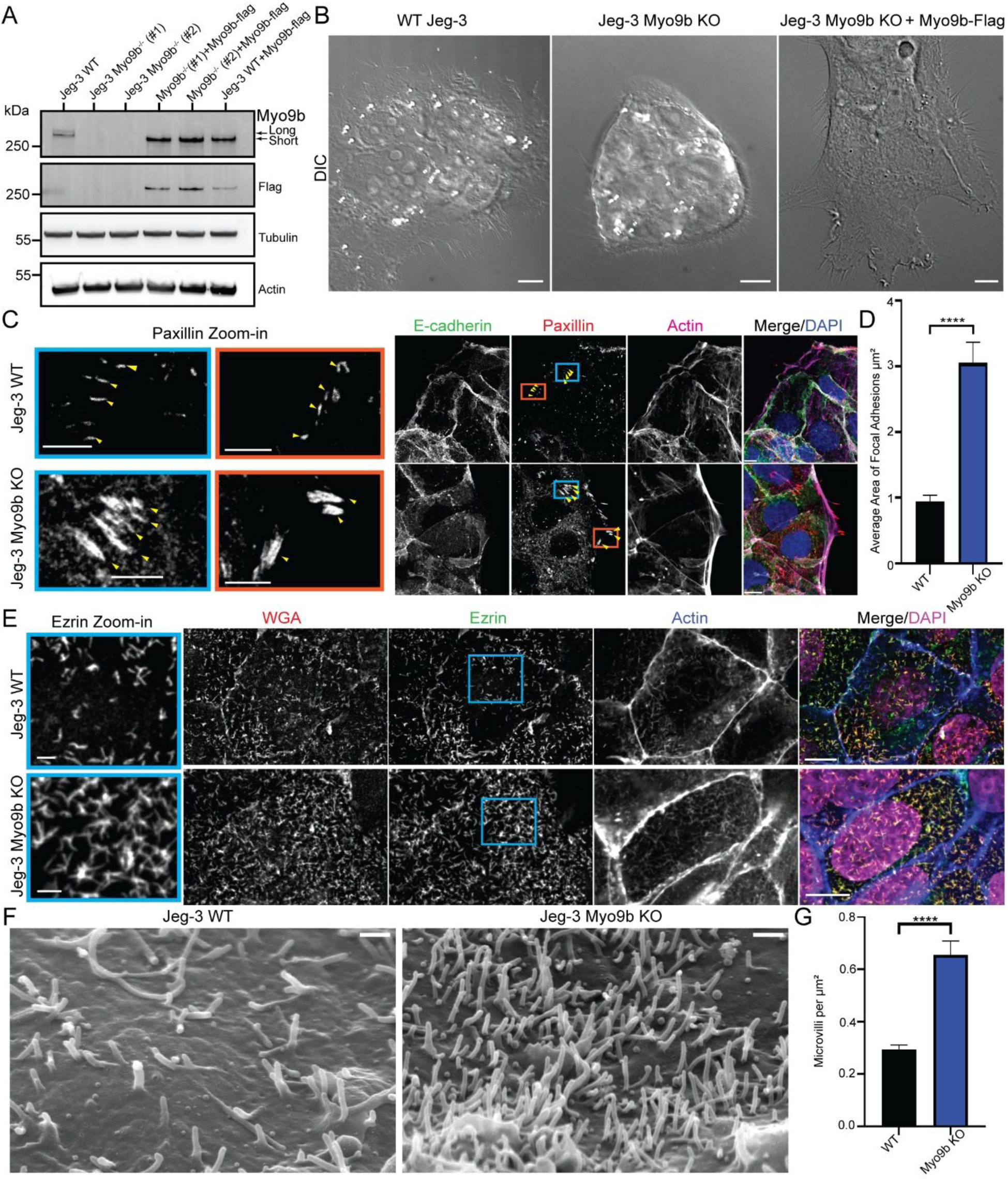
Myo9b^-/-^ (KO) Results in Larger Focal Adhesions and Increased Microvilli Density. A) Western blot identifying Myo9b knockout clones and validation of the antibody targeting human Myo9b. Blot stains against endogenous anti-Myo9b, anti-Flag-tagged Myo9b, anti-tubulin, and anti-beta-actin in Jeg-3 WT cells, Jeg-3 Myo9b KO cells, Jeg-3 Myo9b KO cells rescued with Myo9b-Flag, and Jeg-3 WT cells overexpressing Myo9b-Flag. Long and short indicate the two isoforms of Myo9b endogenously produced through splice variations. Expressed Flag-tagged Myo9b sequence matches the short canonical isoform. B) Differential Interference Contrast (DIC) microscopy of WT, Myo9b KO, and Myo9b KO rescued with Myo9b-Flag. Scale bar 10 μm. C) Spinning disk confocal immunofluorescent imaging of Jeg-3 WT and Myo9b KO cells stained with anti-E-cadherin, anti-paxillin, phalloidin (actin), and DAPI, with zoom in on the boxed area for paxillin staining showing larger focal adhesions in Myo9b KO cells. Scale bar 10 μm; inset 5 μm. D) Quantification of the area of focal adhesions in WT and Myo9b KO cells. Bars represent mean ± SEM; Significance by unpaired t-test, (n = 10 cells each condition, **** =p ≤ 0.0001). E) Spinning disk confocal immunofluorescent imaging of Jeg-3 WT and Myo9b KO cells stained with WGA, anti-ezrin, phalloidin (actin), and DAPI, with zoom in on the boxed area for ezrin staining showing an increase in microvilli density. Scale bar 10 μm; inset 2 μm. F) Scanning electron microscopy (SEM) images of Jeg-3 WT and Myo9b KO cells. Scale bar 1 μm. G) Quantification of the microvilli density on the surface of Jeg-3 WT and Myo9b KO cells from immunofluorescence images. Bars represent mean ± SEM; Significance by unpaired t-test, (n= 30 cells each condition, ****= p ≤ 0.0001).

To characterize the basic morphology of Jeg-3 Myo9b KO cells. Jeg-3 WT, Myo9b KO, and Myo9b KO cells rescued by expression of Myo9b-Flag were imaged using differential interference contrast (DIC) microscopy. The Myo9b KO cells formed domed, tightly packed colonies compared to WT Jeg-3 cells, and this phenotype was rescued by the reintroduction of Myo9b (Fig. 2B). We hypothesized that morphological defects may arise from the loss of Myo9b-specific RhoGAP activity affecting focal adhesions at the leading edge or cell-cell junctions, as tightly controlled RhoA is necessary for their proper formation.(20, 21, 23, 28, 29). Using spinning disk confocal microscopy, we stained Jeg-3 WT and Myo9b KO cells for E-cadherin (adherence junctions), paxillin (focal adhesions), and actin (Fig. 2C). Alterations to E-cadherin were undetectable between conditions. However, paxillin staining revealed a significant increase in the area of focal adhesions in the Myo9b KO cells (Fig. 2C, inset, and D). Additionally, there was an extensive actin bundling phenotype in which actin filaments running parallel to the leading-edge membrane were greatly enriched as measured by actin bundle area quantification (Fig. S2). These focal adhesion and leading-edge actin bundling phenotypes are consistent with previous reports characterizing the functional loss of Myo9b (20, 23).

### Cells Lacking Myo9b Have Increased Microvilli Density and Reduced Microvilli Turnover

Following validation of the Myo9b KO cells and characterization of leading-edge defects, we sought to characterize Myo9b’s function at the apical membrane and microvilli. Immunofluorescent confocal imaging of Jeg-3 WT and Myo9b KO cells stained with non-specific plasma membrane marker wheat germ agglutinin (WGA), the microvilli-specific protein ezrin, and actin, revealed a significant increase in microvillar density upon loss of Myo9b compared to WT cells (Fig. 2E, inset; F, G). Quantification of the ezrin stained microvilli indicated that Myo9b KO cells had more microvilli per square micron compared to WT cells cultured under identical conditions (Fig. 2E, inset, F, G). Reintroduction of Myo9b-Flag to Myo9b knockout cells resulted in the localization of Myo9b-Flag to the tips of microvilli and a restoration of microvilli density to WT levels (Fig. 3A, inset and B). We concluded that Myo9b regulated the abundance of microvilli expressed at the apical plasma membrane.

**Figure 3.**
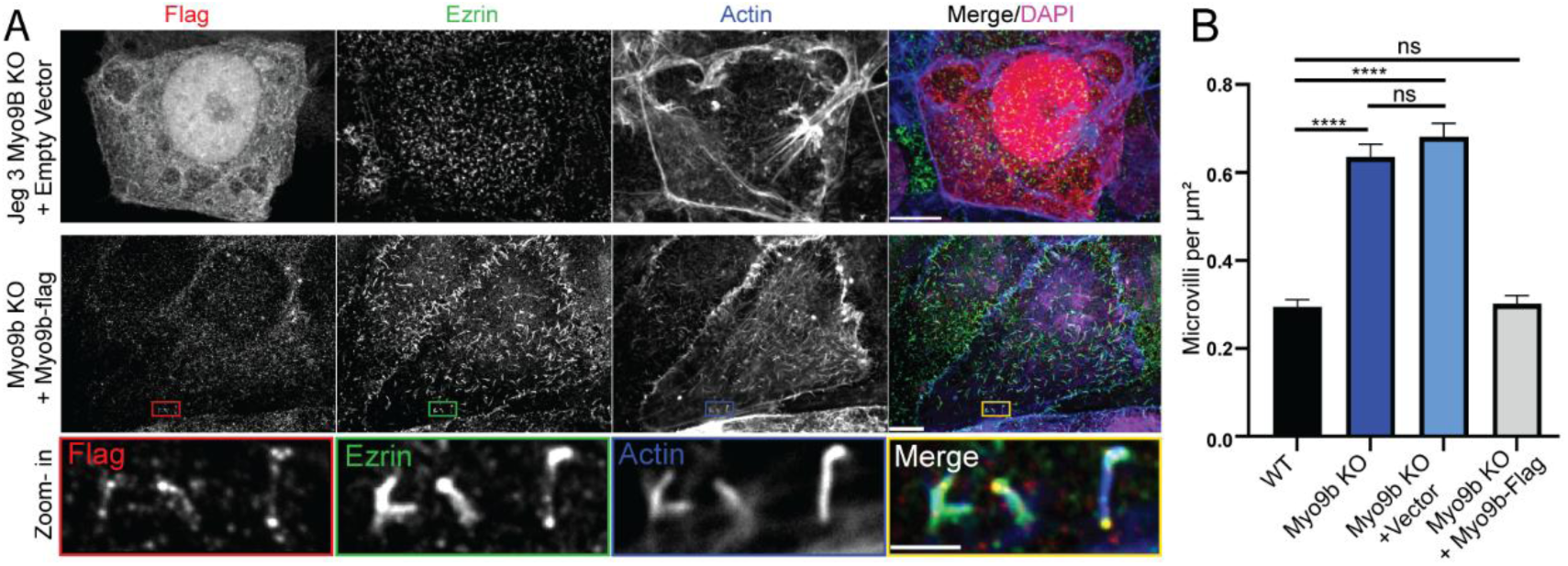
Myo9b KO Microvilli Phenotype Can Be Rescued Along with Localization A) Spinning disk confocal SORA immunofluorescence imaging of Jeg-3 Myo9b KO cells rescued with empty vector-Flag or Myo9b-Flag stained for anti-Flag, anti-ezrin, phalloidin (actin), and Dapi. Zoom-in of Myo9b KO rescued with Myo9b-Flag to show Myo9b-Flag localization at tips of microvilli. Scale bar 10 μm: inset 2 μm. B) Quantification of microvilli density normalized to the cell area. Jeg-3 WT and Myo9b KO conditions were duplicated from Figure 2G for comparison. Bars represent mean ± SEM; Significance by unpaired t-test (n= 30 cells for each condition, **** =p ≤ 0.0001, ns = not significant).

A possibility that could contribute to the increase in microvilli density is a change in microvilli turnover. To determine if Myo9b could regulate microvilli lifetime, we investigated microvillar turnover under live-cell conditions. We expressed eGFP-EBP50, which we had previously used to characterize microvilli lifetime in Jeg-3 WT and Myo9b KO cells, and tracked the persistence of individual microvilli over time (30-32) (Movie 2 and 3, Fig. S3). Quantification of the microvilli lifetimes indicated that microvilli in Myo9b KO cells had microvilli that persisted for significantly longer than in WT cells (Fig. S3). Therefore, Myo9b is necessary for regulating microvillar dynamics, however, the mechanism by which Myo9b altered microvilli abundance and turnover remained unknown.

### The Motor and RhoGAP Domain of Myo9b are Both Required for Regulating Microvilli

Myo9b contains at least two domains that could be responsible for regulating the apical actin cytoskeleton: its RhoA GAP and myosin motor domain. To test whether each domain is required to regulate the apical actin cytoskeleton, we reintroduced GAP or motor-domain variants of Myo9b into Jeg-3 Myo9b KO cells, replacing the endogenous GAP or motor with the mutant variants. Using confocal imaging colocalizing the variants with ezrin and actin, we observed their effects on microvillar organization in addition to the characterization of the motor’s cellular localization and actin organization (Fig. 4A). We mutated the residue 1735 arginine finger in human Myo9b, essential for RhoA-GAP activity, and expressed the tagged Myo9b(R1735M)-Flag (GAP-dead) variant in Myo9b KO cells (10, 11). Expression of the GAP-dead variant resulted in a significant decrease in microvilli density relative to the KO phenotype but did not restore microvilli to WT levels (Fig. 4B, C). These data indicate that the RhoA-GAP domain is necessary for proper microvillar organization. However, the partial rescue indicated that other domains or functions independent of the RhoA-GAP activity of Myo9b also influence microvilli density.

**Figure 4.**
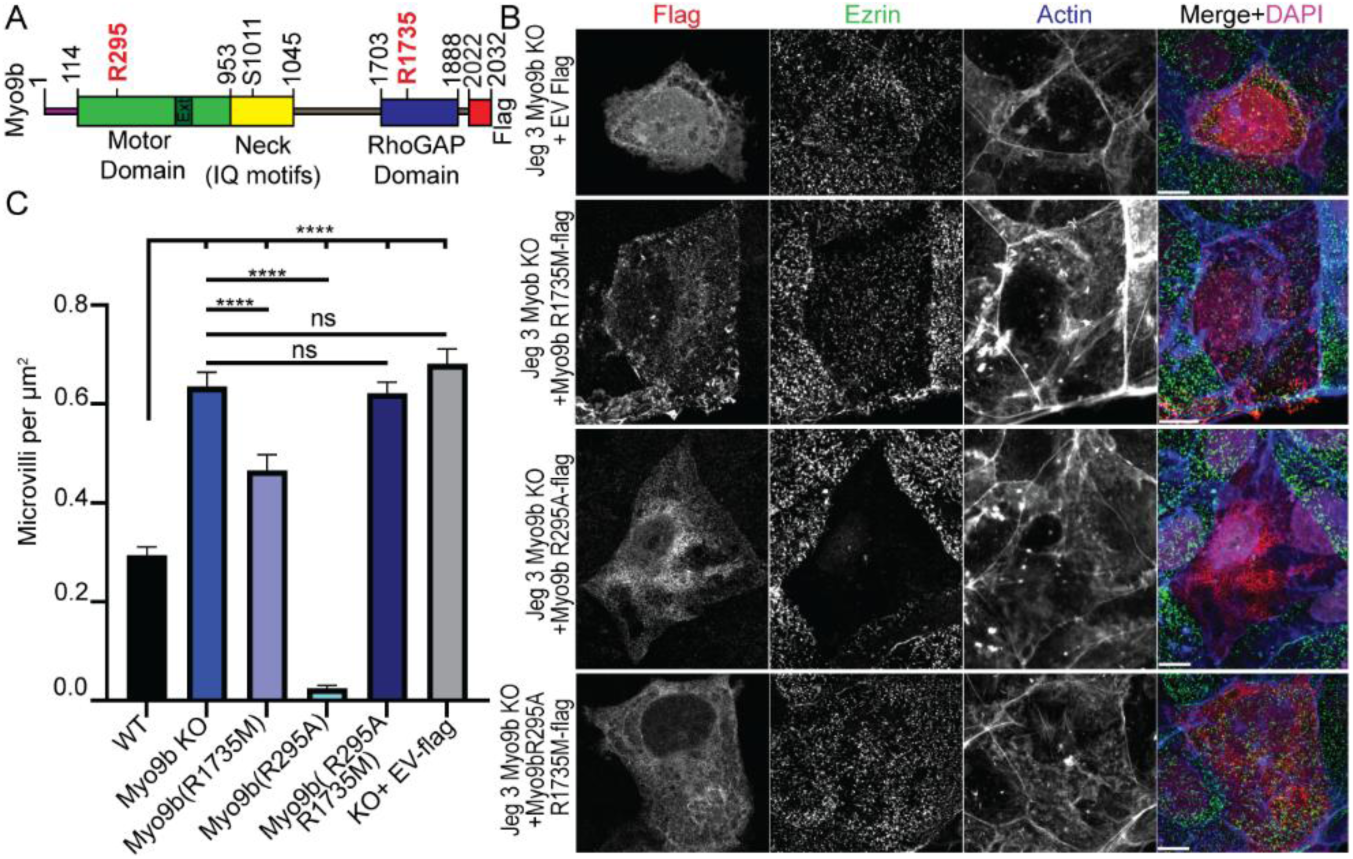
Myo9b’s Motor and GAP Domains are Necessary for Microvilli Regulation. A) Myo9b protein domain schematic. Red text indicates specific point mutations used in Figure 4. B) Spinning disk confocal SORA immunofluorescent imaging of Jeg-3 Myo9b KO cells expressing Flag-tagged empty vector, Myo9b(R1735M), Myo9b(R295A), and Myo9b(R295A and R1735M), stained with anti-flag, anti-ezrin, phalloidin (actin), and DAPI. Scale bar 10 μm. C) Quantification of microvilli density from conditions in (B). Myo9b R295A ablates most microvilli. Microvilli counts were normalized to cell area, and Jeg-3 WT and Myo9b KO conditions were duplicated from Figure 2G, and Myo9b KO +EV-Flag was duplicated from 3D for comparison. Bars represent mean ± SEM; Significance by unpaired t-test (n= 30 cells per condition, **** =p ≤ 0.0001, ns = not significant).

We next tested if the motor domain was necessary for microvilli regulation by Myo9b. We designed a Myo9b(R295A)-Flag (motor-dead) variant with a point mutation ATPase switch-2 salt bridge preventing the ATPase activity of the motor domain (8, 33). This mutation prevents canonical actin binding and force production precluding the possibility for the monomeric motor to crosslink actin. Expression of the motor dead variant resulted in a dramatic loss of microvilli (Fig. 4B, C). In contrast to the native and GAP-dead Myo9b, the motor-dead construct failed to localize within cells and instead was diffusely found throughout the cytoplasm (Fig. 4B). We concluded that the motor domain was necessary for the microvilli-specific localization of Myo9b and that loss of this specificity was catastrophic to the expression of microvilli.

We hypothesized that the motor domain was responsible for localizing the GAP domain to the tips of microvilli, where both the RhoA-GAP domain and other functions of the protein coordinate proper microvilli assembly and maintenance. To test whether the mislocalization of the GAP domain caused the ablation of the microvilli observed in the motor-dead construct, we expressed a full-length Myo9b construct with both motor and GAP mutations, Myo9b (R295A, R1735M)-Flag. The motor-GAP dead construct was localized diffusely throughout the cytoplasm. However, the increased microvilli phenotype persisted, and microvilli density was not significantly different from that in the KO cells (Fig. 4B, C). These data suggest that localization of Myo9b’s RhoA GAP domain to microvilli through the ATPase activity of the motor domain is required for proper microvilli expression.

### Myo9b is Motile in Jeg-3 Cells

Our data indicated that the motor domain of Myo9b was necessary for microvilli tip localization. However, the motility of Myo9b has never been observed in cells to our knowledge. Additionally, Myo9b is reported to have a relatively slow motility rate of only 19.6 ± 7 nm/s along actin *in vitro* compared to most transport myosins, raising the question of whether its motility would be possible to overcome actin treadmilling of the microvilli core bundle (12, 34-36). To investigate whether Myo9b motility could be observed directly within microvilli, a fluorescently tagged StayGold (SG-Myo9b) was expressed in Myo9B KO cells. Using live-cell spinning disk SORA confocal microscopy we were able to achieve a temporal resolution of <200ms and a spatial resolution of ∼150nm enabling us to track resolvable Myo9b motile particles before they accumulated at the tips of microvilli (Movie 4, Fig. 5A). Surprisingly, during imaging in live cells, we observed SG-Myo9b moving at an average speed of 223 ± 117 nm/s (Movie 4 and Fig. 5A, B) approximately a full order of magnitude faster than the measured *in vitro* single molecule monomeric Myo9b velocity (12). To test if the motility was dependent on the ATPase activity of Myo9b we repeated the experiment with the motor dead SG-Myo9b(R295A) variant. When the SG-Myo9b(R295A) construct was expressed in Myo9b KO cells, the fluorescence was diffuse without concentration at the tips of projections, and no directed motility was observed (Movie 5, Fig. 5A).

**Figure 5.**
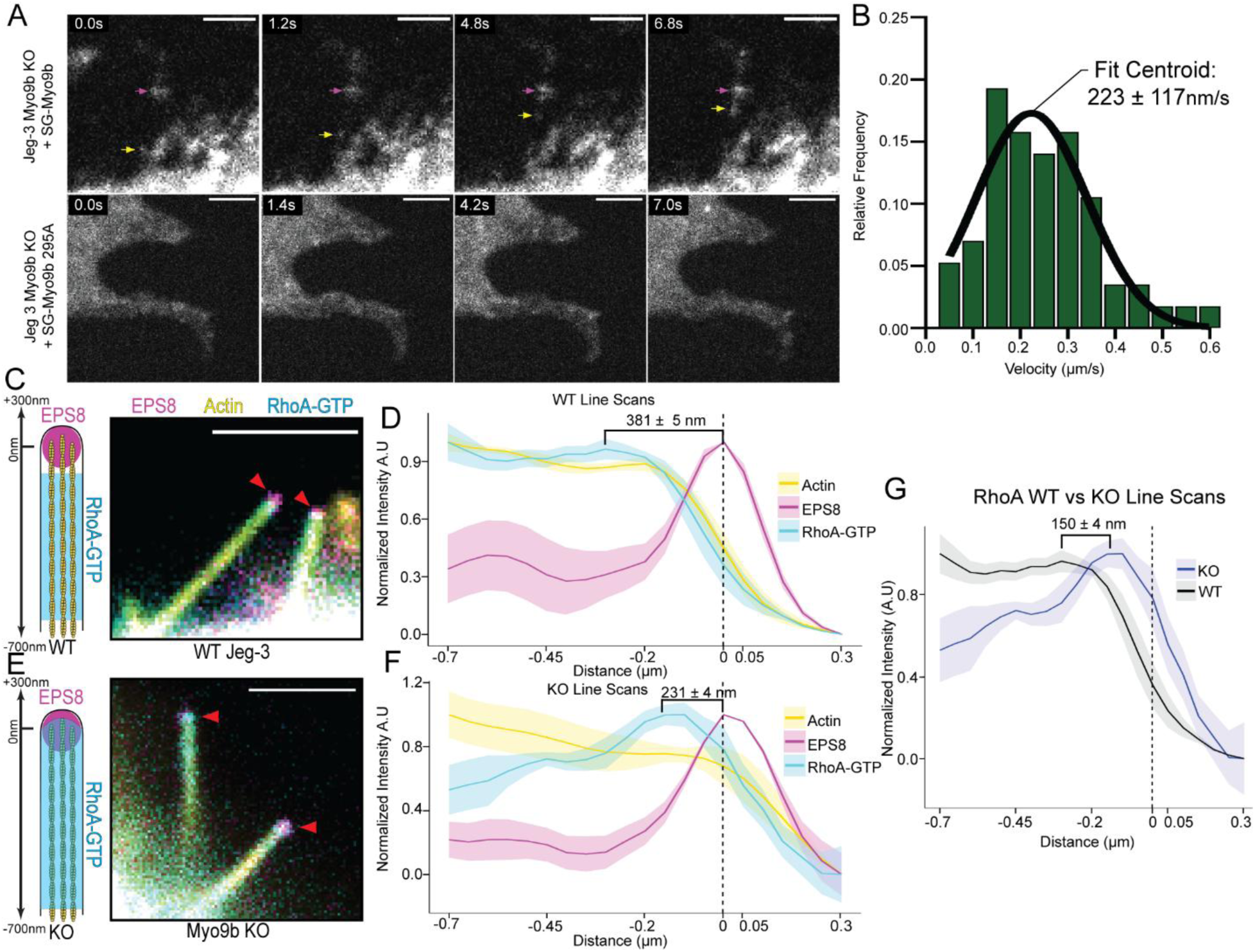
Myo9b is Motile in Jeg-3 Cells and Locally Regulates RhoA. A) Still images from Video 3 showing SG-Myo9b moving up an actin projection (yellow arrows) towards the tip of the projection (magenta arrows) and Myo9b KO + SG-Myo9b R295A. Scale bar 2 μm. B) Histogram and Gaussian fit showing the distribution of SG-Myo9b velocities. C) Schematic showing representative WT microvilli, and still images of microvilli from live cell spinning disk confocal SORA imaging of Jeg-3 WT cells stained with ceurilian-EPS8, dimericTomato-2xrGBD (Active RhoA-GTP biosensor), and SG-Ftractin (actin). Scale bar 2 μm. D) Normalized fluorescence intensity profiles of actin, EPS8, and RhoA-GTP from (C). Intensity profiles were normalized to the EPS8 peak and extend from 700nm before the peak to 300nm after the peak. The distance between the average RhoA-GTP peak and the average EPS8 peak is 381 ± 5 nm. n= 5 cells. E) Representative schematic for KO microvilli, and still images of microvilli from live cell spinning disk confocal SORA imaging of Jeg-3 Myo9b KO cells stained with ceurilian-EPS8, dimericTomato-2xrGBD, and SG-Ftractin (actin). Scale bar 2 μm. F) Normalized fluorescence intensity profiles of actin, EPS8, and RhoA-GTP from (E). The distance between the average RhoA-GTP peak and EPS8 peak for KO is 231 ± 4 nm. n= 5 cells. G) Overlays of the RhoA-GTP intensity of Jeg-3 WT and Myo9b KO from (D) and (F). Distance between average RhoA-GTP peak of WT and KO is 150 ± 4 nm. n=10 cells.

Relative fluorescent intensity using standardized intensity calibration beads revealed that the motile WT SG-Myo9b particles were, on average 6.26±0.27 (mean±SEM) fold higher intensity than the diffusive-like signal observed in the SG-Myo9b(R295A) (Fig. S4). Relative intensity of the stable, tip-localized WT SG-Myo9 measured 15.76±1.9 times the diffusive-like signal, indicating that both the microvilli tip and motile events moving toward the tips were potential oligomers of multiple SG-Myo9b complexed through an unknown mechanism (Fig. S4). Therefore, we concluded that the Myo9b motor domain’s ATP hydrolysis cycle is necessary for the localization of Myo9b at the tips of membrane projections and cellular motility.

### Myo9b Locally Regulates RhoA at the Tips of Microvilli

Our results indicated that proper localization of the RhoA-GAP activity of Myo9b to the tips of microvilli is required for proper microvilli regulation. However, we wanted to directly measure how Myo9b was regulating RhoA in microvilli. To visualize and quantify RhoA activity, we used spinning disk confocal SORA live cell microscopy and expressed an active RhoA biosensor (tdTomato-2xGBD), which preferentially binds the active GTP bound form of RhoA (37). To orient the biosensor between the base, tip, and core sections of the microvilli, we co-expressed cerulean3-EPS8 and F-tractin-StayGold (Actin-SG) into both Jeg-3 WT and Myo9b KO cells (38, 39).

We hypothesized that because Myo9b localizes to the tips of microvilli, the active fraction of GTP-RhoA would be reduced at the tips of microvilli in WT cells. Additionally, we hypothesized that the loss of Myo9b’s GAP activity from the tips of microvilli would result in an increase in active RhoA at microvilli tips in Myo9b KO cells.

To quantify the fluorescent intensity of active RhoA in the microvilli of both WT and KO cells, we performed a line scan of each microvillus tracked over time in our live-cell imaging. Intensity profiles were oriented to the EPS8 peak, which we defined as the microvillus tip. Using the EPS8 peak as a reference point, we aligned all measured microvilli actin and active RhoA biosensor intensity signals from 700nm before the peak to 300nm after the peak for each condition.

In WT cells, the active RhoA fluorescence intensity tapered off just before the EPS8 peak (Fig. 5C, D). In contrast, in Myo9b KO cells, the active RhoA fluorescence intensity peak overlapped with the EPS8 peak (Fig. 5E, F). To directly compare active RhoA signals in WT vs Myo9b KO cells, we set the EPS8 peaks to zero in both conditions and overlaid the active RhoA peaks (Fig. 5G). This analysis indicated that in cells lacking Myo9b, active RhoA fluorescence at the tip of the microvilli was increased compared to WT cells. These data show that Myo9b locally regulates RhoA at the tips of microvilli.

### Myo9b Disease Mutation Mislocalizes Myo9b

Multiple studies have identified a single-nucleotide polymorphism (SNP rs1545620) in the *MYO9B* gene in individuals with intestinal barrier diseases (4-6). This SNP results in expression of the Myo9b point mutant, S1011A, located in the third IQ motif of Myo9b (Fig. 6A). To probe the effect of this mutation, we generated a Flag-tagged Myo9b construct containing the S1011A mutation (Myo9b-S1011A) and expressed it as the only genetic copy in Jeg-3 Myo9b KO cells (Fig. 6A). Expression of Myo9b-S1011A-Flag resulted in a nearly complete ablation of microvilli (Fig. 6B, C). However, unlike either the wildtype or motor dead Myo9b(R295A) constructs, the Myo9b S1011A localized preferentially to the junctions but, like Myo9b(R295A), caused a lack of microvilli (Fig. 6B).

**Figure 6.**
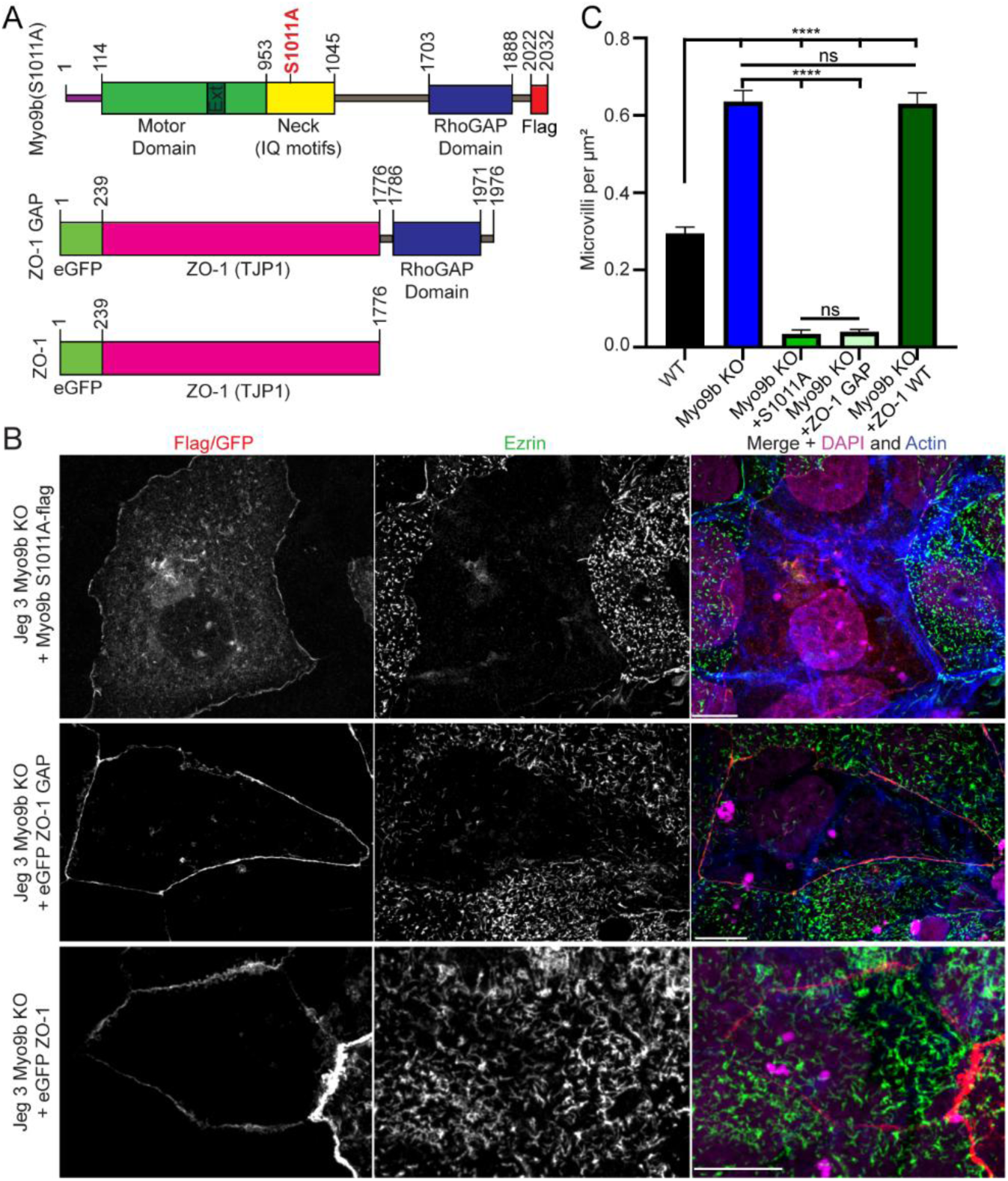
Myo9b Disease Mutation (S1011A) Causes Inappropriate Localization of Myo9b. A) Myo9b disease protein domain schematic. Red text indicates specific point mutations found in the disease state. ZO-1 GAP protein domain schematic. GAP domain is the GAP domain of Myo9b. ZO-1 eGFP control protein domain schematic. All these constructs are used in Figure 6B. B) Spinning disk confocal immunofluorescent imaging of Jeg-3 Myo9b KO cells transiently expressing Myo9b S1011A-Flag, eGFP ZO-1 GAP, or eGFP ZO-1, stained with Flag/GFP, ezrin, actin, and DRAQ5 (DNA). Scale bar 10 μm. C) Quantification of microvilli density from (B). Microvilli counts were standardized to cell area. WT and KO conditions were duplicated from Fig 2G. Bars represent mean ± SEM; Significance by unpaired t-test (n= 30 cells, **** =p ≤ 0.0001, ns = not significant).

To test if the loss of microvilli was due to the inappropriate localization of the GAP domain, we created a construct to generate a synthetic protein containing eGFP-ZO-1 (AKA,TJP1:Tight Junction Protein-1) linked with the GAP domain of Myo9b (Fig. 6A). Expression of the ZO-1 GAP construct in Myo9b KO cells resulted in a localization of the synthetic protein to junctions and a loss of microvilli (Fig. 6B, C). Therefore, we concluded that the S1011A variant disrupts the proper targeting of the Myo9b GAP domain, resulting in severe defects in microvillar biogenesis or maintenance in human cells.

## Discussion

### The Myo9b S1011A Disease Variant Destroys Microvilli

Celiac disease is a multifaceted disorder where no single gene is either necessary or sufficient for development (40). In addition to mutations leading to an aberrant autoimmune response, many IBDs involve an exacerbating mutation that causes a barrier defect in the epithelial layer of the primary gut tissue (1). When epithelial barrier function fails, gut contents inappropriately cross into the underlying cellular matrix or bloodstream. Significant work has advanced the understanding of Myo9b’s influence on leukocyte motility, where Myo9b localizes to the leading edge of motile cells, regulating actin polymerization (21, 22, 41). However, one conundrum has been that despite numerous bioinformatic analyses linking the S1011A mutation to IBD, mice lacking the Myo9b gene exhibit defects in immune cell motility (19, 20) and abnormal intestinal barrier function (42), do not develop any overt autoimmune disease.

Our study is the first to characterize a potential disease mechanism for the Myo9b S1011A variant in human epithelial cells involving the regulation of apical cell morphology and junctions. Our data indicate that under normal conditions Myo9b is confined to a nanoscale localization at the tips of microvilli, in part to regulate RhoA signaling and actin polymerization. However, in cells where the S1011A variant is the only Myo9b copy, the protein is inappropriately localized to the junctions, resulting in a near total loss of microvilli (Fig. 6B, C). Shockingly, this disease variant localization is potentially worse than the genetic loss of Myo9b where cells lacking Myo9b express ample microvilli, albeit in a dysregulated turnover state. The loss of microvilli in the S1011A variant condition appears to coincide with the mislocalization of the RhoA-GAP domain as targeting just the GAP domain to tight junctions using Zo-1 had the same result (Fig. 6B, C).

These data suggest a novel finding that Myo9b disease mutants may not represent a loss of function but instead gain the ability to specifically target junctions. Serine 1011 is positioned within the third IQ calmodulin-binding motif of Myo9b, placing it at a critical sequence for motor and force production regulation (43). The mechanism underlying the potential loss of calmodulin/myosin light chain binding at the third IQ motif in the S1011A variant that results in mislocalization to junctions will require further investigation. Further, our *in vivo* data suggest that Myo9b may also be enriched at the base of microvilli in the terminal web (Fig. 1C-E) of actin which will require additional study.

### Myo9b’s Domain Structure Enables Multiple Mechanisms for Cytoskeletal Regulation

Myo9b possesses at least two domains that directly regulate the apical actin cytoskeleton: its motor domain (12, 13, 44) and its RhoGAP domain (10, 18). Previous studies have focused on these two domains, describing Myo9b as a ‘motorized GAP’ (7, 8, 21). However, the GAP-dead variant of Myo9b partially rescues the KO Myo9b microvilli phenotype, indicating that the GAP domain itself is not the sole regulator of apical actin (Fig. 4B). One explanation is that in microvilli, Myo9b may also use its motor domain to regulate actin filament organization. The loop 2 extension found in the human isoform Myo9a confers a second actin-binding site to cross-link parallel actin filaments (45).

Yet other uncharacterized potential mechanisms exist within Myo9b, including a N-terminal extension of the motor domain containing a potential Ras-association domain and a Phorbol-ester/DAG-type C1 domain between the IQ motifs and the GAP domain, both of unknown relevance. Our data supports a model where Myo9b uses both its RhoA GAP domain and its motor domain to locally regulate the apical actin cytoskeleton. Yet our data also raises new questions as to the functions of additional structural features of Myo9b. Specifically, is the potential lipid associated DAG-type C1 domain necessary for localization to the apical plasma membrane and what is its role in our observed motility within cells.

### Myo9b’s Cellular Motility is Faster than the Purified Single Motor Velocity

The continuous recycling of actin monomers within the bundles of microvilli core filaments (i.e., actin treadmilling) could be a limitation for a relatively slow motor Myo9b to localize to the tips of projections. Depending on actin assembly conditions within the microvillus, actin treadmilling ranges from about 3nm/s to 21nm/s (34-36) Specifically, *in vitro* Myo9b has been predicted to move at 19.6 ± 7 nm/s (12), slower than or equal to the rate of actin treadmilling. Our live-cell imaging data on SG-Myo9b, showing Myo9b moving within cellular projections at approximately 223 ± 117 nm/s, would overcome this issue. The loss of Myo9b’s ATPase activity using a motor dead (R295A) variant kills all motility and Myo9b localization (Fig. 4B). Interestingly, these data present multiple models for the mechanism of Myo9b motility in cells.

The most straightforward interpretation is that Myo9b forms a processive complex with its activating or binding partners that move along actin at higher velocities than the purified *in vitro* monomeric motor alone. This model has been identified in numerous other cytoskeletal motors (46, 47); however, crosslinking pulldowns identified Myo9b as a monomeric motor (13). An additional possibility is that Myo9b is cargo for another myosin found in microvilli, such as Myosin V, Myosin VII, or Myosin XV (48). This mechanism has been previously observed in microtubule-based motors where inactive motors “hitchhike” with cargos transported by other motor types (49, 50). Under the hitchhiking model, the motor domain R295 residue and resulting ATPase activity may be required for the activation of Myo9b and for structural unfolding from an autoinhibited state to bind to the motile complex. Nonetheless, it is clear that the ATPase activity of Myo9b is necessary for the motor’s movement toward the tips of actin protrusions. Reconstitution or isolation of the complete cellular motile complex will be required to answer how Myo9b achieves our measured cellular velocity.

## Materials and Methods

### Reagents and cDNAs

The open reading frame (ORF) Accession number NM_001130065.2, Human MYO9B cDNA, was obtained from Genescript (Piscataway, NJ) and verified by whole plasmid sequencing. Myo9b molecular biology constructs were generated using polymerase chain reaction (PCR) with KOD1 polymerase (Sigma 71085-3) or PrimeSTAR GXL polymerase (Takara-Bio R050A). MYO9B variants were produced by site-directed mutagenesis using the same polymerases. PCR and DNA cleanup kits (NEB T1030S) and GeneJET Gel Extraction kits (ThermoFisher K0692) were used for purification of PCR products and for InFusion HD cloning (Takara-Bio 639650) of genes into mammalian expression vectors. Constructs were transformed into chemically competent DH5alpha (ThermoFisher C404006), STBL3 (New England Biolabs C3040I, or Stellar (Takara-Bio 636766) bacteria, selected for antibiotic resistance, and whole plasmid sequence verified. pcDNA3/mStayGold(c4)=UtrCH was a gift from Atsushi Miyawaki (Addgene plasmid # 212020; http://n2t.net/addgene:212020; RRID:Addgene_212020) (39). The UtrCH gene was exchanged with human WT Myo9B for N-terminal tagging with mStayGold via a c4 adaptor and a linker [(GGGGS)3] using NEB Gibson Assembly (Cat #E5510S). Constructs were then transformed into STBL3 chemically competent cells. Using ampicillin resistance successfully cloned constructs were selected and whole plasmid sequenced verified.

The mCerulean3-C1 was a gift from Michael Davidson (Addgene plasmid # 54605) (51) mCerulean3-C1-Eps8 was constructed by inserting the PCR-amplified Eps8 sequence at the EcoRI site of mCerulean3-C1using InFusion HD cloning (Takara-Bio 639650). The F-tractin=mStayGold was a gift from Atsushi Miyawaki (Addgene plasmid # 21201) (39) and tdTomato-2xrGBD was a gift from Dorus Gadella (Addgene plasmid # 176098) (52). The eGFP ZO1 Gap plasmid was constructed by inserting a SpeI site at the C-terminus of ZO1 in the ZO1-eGFP plasmid (Addgene 30313) using site-directed mutagenesis. The GAP domain of Myo9B was amplified by PCR and cloned into the SpeI site by InFusion HD cloning (Takara-Bio 639650).

### Cell Culture

The Jeg-3 cells (ATCC HTB-36), LLC-PK1 Cells (ATCC CL-101), CaCo-2 cells (ATCC HTB-32), DLD-1 colorectal adenocarcinoma cell lines (ATCC HTB-37 and ATCC CCL-221), and HeLa (ATCC CRM-CCL-2) cell lines were obtained commercially through ATCC.org with approval under the University at Buffalo protocol (030BIO00000293). The Jeg-3 cells were maintained in MEM (Thermo Fisher Scientific; Cat# 10370-021) supplemented with 10% fetal bovine serum (FBS; ThermoFisher Scientific; Cat# 26140079) 10% GlutaMAX (ThermoFisher Scientific Cat# 35050061) and 100mg/mL penicillin/streptomycin (ThermoFisher Scientific; Cat# 15070063). The PK1 and Hela cell lines were maintained in DMEM (Thermo Fisher Scientific; Cat# 11965-092) supplemented with 10% FBS (ThermoFisher 26140079), 10% GlutaMAX (ThermoFisher 350500-61), 1mM sodium pyruvate (Thermo Fisher Scientific; Cat# 11360-070), and 100mg/mL penicillin/streptomycin (ThermoFisher 15070063). The DLD-1 cells were grown in RPMI 1640 (Corning: Cat# 10-040-CV) supplemented with 10% FBS (ThermoFisher 26140079), 10% GlutaMAX (ThermoFisher 350500-61), and 100mg/mL penicillin/streptomycin (ThermoFisher 15070063). All cell lines were maintained in 100 mm TC-treated culture dishes (Corning 430167) in a humidified incubator at 37 °C and 5% CO2. Cells were transiently transfected into mammalian cells with PEI Max (Polysciences 24765).

### CRISPR Myo9b Knockout Lines

Benchling Inc (San Francisco, CA) CRISPR analysis tools were used to created single-guide RNAs (sgRNAs). The sgRNAs were cloned into puromycin resistant pLenti-CRISPRV2 (Addgene; Cat# 49535) described previously (53). Three separate Myo9b KO lineages were created using different CRISPR/CAS9 target guides: 5′-GGAGGCATGCTGAAGCCAGG-3′, 5′-GCTGGGGGTAGATGTGCAGG-3′ and 5′-CGGCAGGATGAGTGTGAAAG-3′. Three separate HEK293Tn cell cultures were transfected with the sgRNA sequences against human Myo9b with lentivirus packaging plasmids psPAX3 and pCMV-VSV-G (a gift from Jan Lammerding, Weill Institute for Cell and Molecular Biology, Cornell University, Ithaca, NY). The transfected media containing viral particles was collected after 48 and 72 hours and filtered using a 0.45-micron filter followed by a 0.22-micron filter. Cells were transduced with the filtered virus in media supplemented with 0.8 ug/mL polybrene (Millipore Sigma; #Cat TR-1003-G) and incubated overnight. Viral transduction was repeated twice a day, 7 hours apart for 2 days. After 24 hours, the cells were washed with PBS and transferred to a 100 mm dish containing media supplemented with 2 ug/ml of puromycin. Cells were subjected to single-cell sorting, expanded in puromycin selection and then screened via western blot and immunofluorescence to confirm lack of Myo9b protein expression. The more uniform phenotype expressing was produced by the 5′-CGGCAGGATGAGTGTGAAAG-3′ and this guide was used for experiments where two clones were used to validate the initial phenotypes.

Jeg-3 Myo9b KO cell line was used to create the stable expression cell line of Jeg-3 Myo9b KO + Myo9b flag. Similar to above HEK293Tn cells were transfected with psPAX3 and pCMV-VSV-G lentivirus packing plasmids along with the expression vector: pCDH-Blast2.0-Myo9b-Flag using Lipofectamine (Invitrogen; Cat# 13778-030) transfection reagent (for every 5ug of DNA add 10uL of lipofectamine) for lentiviral particle production. The media containing viral particles was collected and cells were transduced following the same methods as above. Cells were then single cell sorted and expanded in 10 ug/mL Blasticidin. The expression of Myo9b-flag in the sorted cells was confirmed using Western blot analysis and immunofluorescence.

### Myo9b Antibody

The antigen against human Myo9b was produced by expressing a truncated region of Myo9b (AA 1054 to 1484) identified as unique using protein BLAST and specifically maximizing the distinct sequnce from human Myo9A. The Myo9b-fragment was expressed linked to an N-terminal-SUMO-HIS-tagged in bacteria and purified using NiNTA resin (QIAGEN; Cat# 1018244). Purified ULP1 was used to cleave the SUMO tag creating a untagged Myo9b fragment, and the protein was passed through a HiLoad 16/600 Superdex 200 size-exclusion chromatography column (Cytiva 28989335) mounted on an FPLC (General Electric AKTA pure). Antibodies were produced after exchanging the protein into PBS, followed by denaturation by boiling for 5 minutes after the addition of 1 mM DTT (ThermoFisher R086). The protein samples were then snap-frozen in liquid nitrogen and shipped to Pocono Rabbit Farm & Laboratory, Inc (Canadensis, PA) for antibody production in rabbits and further purified by passing the produced sera over an affinity column of the purified antigen. The specificity of the produced affinity purified antibody against Myo9b was confirmed by Western blot (1:1000, see Figure 2A). The animal use protocol was approved by the University at Buffalo (Pocono Rabbit Farm IACUC: PRF2A and Lombardo Lab protocol: ID 030BIO00000293).

### Western Blotting

Cell lysates were resolved using handcast SDS-PAGE 5% or 8% gels and transferred onto a polyvinylidene difluoride (PVDF) membrane using a Transblot Turbo (Bio-Rad 1704150). The membrane was blocked with 5% milk in TBS-T, composed of Tris Buffered Saline (TBS) and 0.5% Tween-20, followed by incubation with primary antibodies diluted in 5% BSA in TBS-T for an hour. Primary antibodies used included: anti-Myo9b (custom-made, 1:1000, rabbit monoclonal), anti-Tubulin (DHSB 12G10, 1:1000, mouse monoclonal), anti-Flag (Sigma, F1804, 1:1000, mouse monoclonal), anti-Actin (Sigma, MAB-1501, 1:1000, mouse monoclonal). Protein bands were detected using fluorescent secondary antibodies including anti-Mouse-680 nm (Invitrogen A21057) and anti-Rabbit-800 nm (Invitrogen A32735). BioRad ChemiDoc was used to image blots and the Western blots were quantified using the intensity profile built-in Gel-Analyzer toolset in ImageJ, and graphs were plotted using GraphPad Prism (GraphPad Software, Inc., Version 11).

### Mouse Intestine Immunofluorescence

Mouse Ilium tissue processing, slide preparation with cryosections, and staining were performed as described by Murray et al. 2026 (54). The dilutions for the primary antibodies were anti-Myosin 9b (custom, 1:100, rabbit monoclonal) and anti-Ezrin (gift from A. Bretscher, 1:100, mouse monoclonal). Secondary antibodies were used at the following dilutions: goat anti-rabbit Alexa Fluor 568 (1:500, Invitrogen; Cat# A10042), goat anti-mouse Alexa Fluor 488 (Company, cat#,1:500), phalloidin 647 (1:500, Invitrogen; Cat# A32723), and Hoechst 33342 (AnaSpec Inc., 83218, 1:3000). Samples were then washed with 1X PBS and mounted with Fluoromount-G (Invitrogen, 00-4958-02). Images were captured with Yokogawa CSU-X1 spinning-disk Marianas microscope (Intelligent Imaging Innovations). All animal protocols and procedures were performed in the laboratory of Christine E. Schaner Tooley and approved by the SUNY Buffalo Animal Care and Use Committee under IACUC protocol BCH08076N.

### Fixed Cell Immunofluorescence

Cells were grown on 1.5 thickness glass coverslips (Warner Instruments; Cat# 64-0734) in 6-well plates. They were washed with PBS three times and fixed with 10% paraformaldehyde (PFA) for 10 minutes. After PBS washes, the cells were permeabilized with 0.2% Triton-X-100, followed by another set of PBS washes. The cells were then blocked with 2% FBS fetal bovine serum (Thermo Fisher Scientific; cat# 26140079) for 30 minutes, followed by incubation with the primary antibody, diluted in 2% FBS, for 1 hour at room temperature. The dilutions for primary antibodies were anti-Myo9b (custom, 1:100, rabbit monoclonal), anti-Ezrin (Gift from A. Bretscher, 1:200, rabbit polyclonal), anti-Flag (Sigma; cat# F1804, 1:250), anti-Paxillin (Abcam; cat# AB 32084, 1:100), and WGA 488 (Invitrogen Cat# W11261, 1:300). The coverslips were washed with PBS three times for 5 minutes each, then incubated with secondary antibodies for 1 hour at room temperature. The secondary antibodies were diluted in 2% FBS, and the dilutions were: anti-Mouse-488 (Invitrogen; cat# A32733, 1:250) and anti-Rabbit-568 (Invitrogen; cat# A10042, 1:200). The nucleus and actin were stained with DAPI (Invitrogen; cat# D1306, 1:10,000) and Phalloidin-647 (Invitrogen; cat# A22287, 1:200) respectively during the secondary antibody incubation. The coverslips were washed three times in PBS for 5 minutes each, mounted using Prolong^TM^ Diamond Antifade Mountant (Invitrogen; Cat# P36961) on microscope slides for imaging (Fisher Scientific; Cat# 12-544-3).

### Confocal Imaging

Fixed cell SORA and confocal immunofluorescent imaging were performed at room temperature using a spinning-disk (Yokogawa CSU-W1) Intelligent Imaging Innovations Marianas microscope with a x63/1.4NA Zeiss objective. The images were captured with a Hamamatsu ORA Quest camera. SlideBook 7 software (Intelligent Imaging Innovations) was used to assemble Z-slices of captured images. Maximum Z-projections were put together in SlideBook 7. For apical surface imaging of anti-ezrin maximum Z projections were generated from Z slices at the apical surface. All maximum Z projections were exported to Adobe Illustrator for editing.

### Live Cell Immunofluorescence

Confocal live cell imaging and SORA live cell imaging of cells expressing GFP-EBP50 (55), SG-Myo9b, SG-Myo9b(R295A), ceurilian-EPS8, dimericTomato-2xrGBD (Active RhoA biosensor), F-tractin=mStayGold, was done on a spinning-disk (Yokogawa CSU-W1) Intelligent Imaging Innovations Marianas microscope with a x63/1.4NA Zeiss objective, Hamamatsu ORCA-Quest camera, SlideBook 7 software (Intelligent Imaging Innovations), and definite focus control. 24 hours before imaging, Jeg-3 cells were transiently transfected, and immediately before imaging, the cell medium was changed to MEM, no glutamine, no phenol red (ThermoFisher Scientific; Cat# 51200-038) supplemented with 10% FBS (Thermo Fisher Scientific; Cat# A56707-01), GlutaMAX (Thermo Fisher Scientific; Cat# 35050-061), and ProLong^TM^ Live Antifade Reagent (1:100) (Thermo Fisher Scientific; Cat# P36974). The cells were then kept in a humidified environmental chamber with 5% CO_2_ at 37°C while imaging. GFP-EBP50-expressing Jeg-3 cells were imaged every 30 seconds for a minimum of 20 minutes to get microvilli turnover. Jeg-3 cells expressing td-Tomato EBP50 and SG-Myo9b were imaged every 0.8 seconds. For Jeg-3 cells expressing ceurilian-EPS8, dimericTomato-2xrGBD, and F-tractin=mStayGold, live cell Z-stack imaging was performed. For live cell motility tracking of cells expressing SG-Myo9b and SG-Myo9b(R295A) cells were imaged every 0.2 seconds. To measure particles that contained motility SlideBook 7 software (Intelligent Imaging Innovations) measurement tool was used to record distance traveled in µm. The start and end frame of motility of was used to calculate duration (seconds).

### Scanning Electron Microscopy (SEM)

The Microscopy Imaging Center at the University of Vermont (Burlington, VT, RRID#:SCR_018821) was used to perform SEM imaging. Cells were cultured on Thermanox coverslips and were fixed at 4°C using Karnovski’s fixative (2.5% glutaraldehyde, 2% paraformaldehyde 0.1M cacodylate buffer, pH 7.2) for 1 h. Then cells were washed 4 times in 0.1 M cacodylate buffer, pH 7.2. For post fixation, cells were treated with 1% osmium tetroxide in 0.1 M cacodylate buffer pH 7.2 at 4°C for 1 h and then rinsed three times. Then, cells were submerged in 1% tannic acid in 0.05 M cacodylate buffer at room temperature for 1 h and washed into 0.05 M cacodylate buffer and then into water. The cells were incubated for 1 h at RT using 0.5% uranyl acetate in MilliQ water and then rinsed into water. The cells were then stored overnight at 4°C in 0.05 M cacodylate buffer. Next, the cells were dehydrated through a graded series of ethanol washes to 100% anhydrous ethanol, followed by critical point drying with liquid CO_2_. Aluminum specimen mounts with a conductive carbon paint were used to mount the samples and then dried overnight using desiccation. The desiccated cells were sputter coated in a polaron sputter coater (Model 5100) with gold/palladium and then stored in dissection conditions. Imaging was performed on the prepared samples using a JSM-6060 scanning electron microscope from JEOL USA, Inc (Peabody, MA).

### Segmentation

Image analysis was performed using Python 3.13.5. For each cell, paired actin and paxillin fluorescence images were loaded, and converted to floating-point arrays. Each channel was intensity-normalized using percentile-based scaling. Images were padded by 60 pixels (px) and then used to define the analyzable cell-edge region which is then removed to prevent image edge from being used as the cell boundary segmentation. Actin and paxillin channels were combined by taking the pixel-wise maximum signal, smoothed with a Gaussian filter, and thresholded to identify low-signal extracellular background. Only background connected to the image border was retained as external background, preventing internal low-signal regions from being misclassified as outside the cell. The cell mask was defined as the complement of this external background and was cleaned by small-object removal, hole filling, and light morphological smoothing.

The cell boundary region was defined as an inward-facing band from the detected external background into the cell mask of 600 px. The external background itself was excluded from all downstream analysis. To better preserve faint true cell edges, the first-pass external background estimate was refined using paired actin and paxillin signal near the cell edge, allowing dim but connected peripheral signal to be reclaimed as cell boundary.

Actin bundles were segmented within the boundary mask from the normalized actin channel after Gaussian smoothing and a sigma of 1.0. Boundary-localized actin signal was thresholded by percentile intensity at the 60th percentile, followed by removal of small objects below the minimum area cutoff of 50 px or 0.125 µm². Focal adhesions were segmented within the same boundary mask from the normalized paxillin channel after Gaussian smoothing with a sigma of 0.85 px and percentile-based thresholding at the 60th percentile as well. Paxillin-positive objects were filtered by a minimum focal adhesion area of 25 px or 0.0625 µm² and a maximum object area of 5000 px or 12.5 µm² to retain focal adhesion-sized structures. Per-cell actin fiber and focal adhesion features were then calculated, including mean actin bundle intensity, total actin bundle area, average actin bundle area, actin intensity normalized by actin bundle area, and actin alignment index, average focal adhesion area, focal adhesion density normalized to boundary area, and average focal adhesion elongation. The area of Paxillin-stained focal adhesions at the cell periphery was quantified using the Area measure tool in ImageJ. Ten Jeg3 WT and Jeg3 Myo9b KO cells were used in the analysis

### Microvilli Quantification and Turnover

To determine microvilli density, cells were stained for anti-ezrin as a marker for microvilli. The number of microvilli in a cell was quantified using the MTrackj plugin in ImageJ developed in (56). The total microvilli number was normalized to cell area to account for cell size differences. The area of each cell was measured by the polygon selection tool in ImageJ. The Microvilli/µm^2^ density values were input into GraphPad Prism in order for statistical analysis. For microvilli turnover, cells were transiently transfected with eGFP-EBP50 a microvilli marker. Then the ImageJ plugin Mtrackj was used to track microvilli lifetimes. For Jeg-3 Myo9b KO cells microvilli at the edge of the cells were only track as the high microvilli density throughout the cell made tracking difficult in the center of the cell.

### Fluorescent Intensity Line Scans and Relative Intensity Analysis

Mouse intestinal tissue line scans cells were rotated to be horizontal in ImageJ. A box using the box tool in ImageJ was drawn from the basal surface of the cell past the apical surface of the cell using the phalloidin (actin) signal. This box was kept consistent for each cell measured to ensure distance from basal surface was kept consistent. The basal surface of the cell was set to 0 µm and fluorescence intensity was measured for each signal (Dapi, anti-ezrin, anti-Myo9b, and phalloidin) using intensity profile toolset in ImageJ. Fluorescent intensity values were input into Excel where a 5 point moving average was applied for each signal in each replicate. Fluorescent intensity for each signal in each replicate was normalized (0-1 A.U.) and then averaged, and SEM was calculated for error. RStudio was then used to graph the normalized average fluorescent intensity ± SEM.

For microvilli fluorescent intensity line scans microvilli were rotated to be horizontal in ImageJ. Then using the Ftractin=mStayGold (actin) fluorescent intensity a box was drawn from the base of the microvilli to past the tip of the microvilli using the box tool in ImageJ. This box was kept consistent across all three signals for an individual microvillus (ceurilian-EPS8, dimericTomato-2xrGBD, and F-tractin=mStayGold). Fluorescent intensity was measured for each signal using ImageJ intensity profile toolset. The fluorescent intensity values were input into Excel where a 3-point moving average was applied. To normalize the size differences across the different microvilli, each replicate was normalized to the EPS8 peak, where for each replicate the EPS8 fluorescent intensity peak signal towards the tip of the microvilli was set to 0 nm and a 1 µm range was selected that covers the peak intensity (700 nm prior to peak and 300 nm after peak). This 1 µm range was applied to the other signals within that replicate. The fluorescent intensities across replicates were averaged and then normalized (0-1 A.U.). The normalized average fluorescent intensity was graphed along with average ± SEM for error in RStudio. Measurement of the distance between intensity peaks in the live cell RhoA-Biosensor traces were calculated by averaging the distance in nm from the EPS8 peak to the RhoA-GTP biosensor peak and error across all replicates. To calculate the difference between the two RhoA-GTP peaks in WT vs. Myo9B KO cells the two averages were subtracted to find their difference relative to the EPS8 peak.

Relative intensity of SG-Myo9b signal at the tips of microvilli, motile events, and diffusive-like events were compared by creating a standardized intensity curve using InSpeck Green (505/515) Microscope Image Intensity Calibration Kit (Thermo-Fisher Cat. # I14785). Beads were mounted in the provided kit mounting media at a 1:10 dilution on 1.5 thickness glass coverslips (Warner Instruments; Cat# 64-0734). Confocal imaging was then performed under identical conditions to the imaging of SG-Myo9B live-cell tracking used for velocity measurements. A standardized curve of 0(no fluorescence), 0.3%, and 1% incorporation of InSpeck Green into the beads was created by measuring the raw intensity of a 5x5 pixel area of individual beads and plotting as a function of the % incorporation. This curve fitted to a linear scale where the slope represents the detected signal as a function of the %incorporation in fluorescent particles while the intercept represented the background fluorescence signal. Intensity from SG-Myo9b-WT was then measured using the same 5x5 pixel raw intensity at the tips of microvilli and the observed motile events while diffusive-like particles were measured in the SG-Myo9b(R295A) motor dead variant condition. Average background subtracted intensities were normalized to the diffusive-like motor dead condition intensity and graphed as a violin-plot using GraphPad Prism.

### Statistical Methods

GraphPad Prism was used to perform statistical analysis. The statistical tested used, the number of independent data points (n), and the calculation of error bars are indicated in the figure legends respective to data analyzed. Nonparametric or parametric statistical analysis was determined based on whether the tested data contained a normal distribution or not.

## Acknowledgments

We thank Jim Sellers, Sarah Heissler, and Joe Cirilo for their helpful discussions. SEM imaging was performed at the Microscopy Imaging Center at the University of Vermont (Burlington, VT, RRID#:SCR_018821). Funding was supported by the National Institutes of Health grant R35GM156870 to Andrew T. Lombardo, grant R01HL163168 to Yongho Bae, and grant R35GM144111 to Christine E. Schaner Tooley. We thank Zack Arthur for editorial review.

