## Supplemental Figures for "Myosin-9b Controls Epithelial Brush Border Architecture through Motility-Dependent RhoA Signaling"

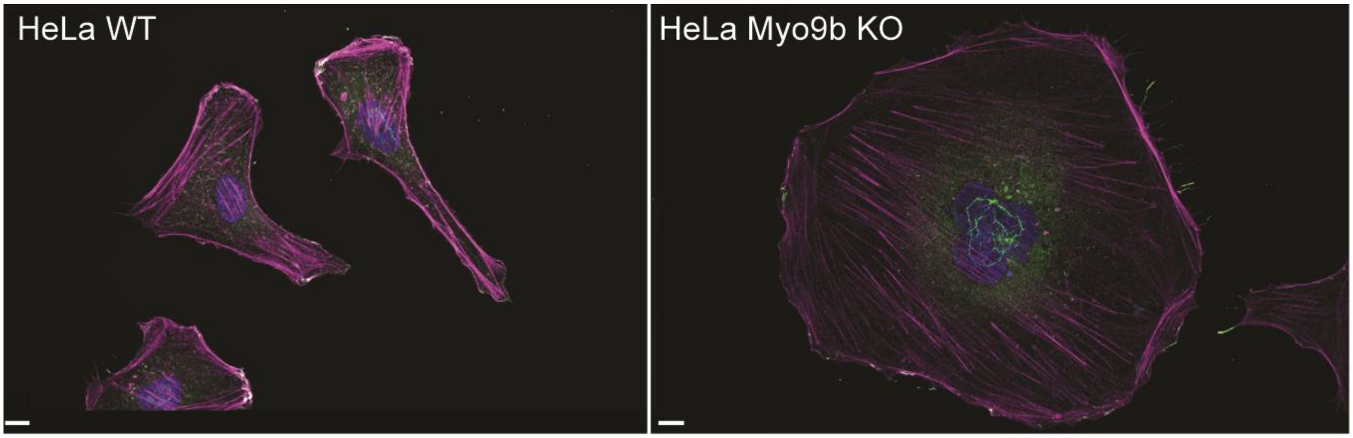


**Fig. S1**. Myo9b KO in HeLa cells results in large multinucleated cells. Spinning disk confocal immunofluorescent imaging of HeLa WT and HeLa Myo9b KO cells stained with DAPI (blue), anti-Ezrin (green), and Phalloidin (Actin- pink). Scale bar 10 μm.

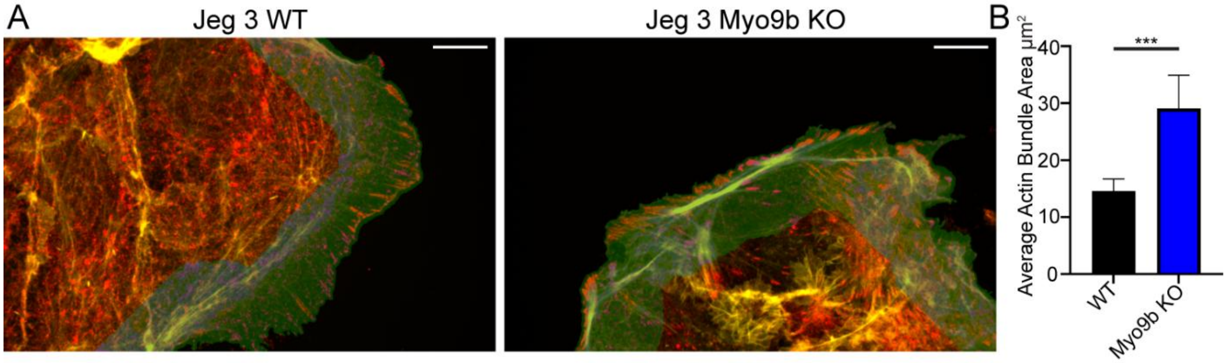


**Fig. S2**. Boundary-localized segmentation of actin bundles and focal adhesions in Jeg3 cells.

A) Representative overlay images generated from paired actin and paxillin spinning disk confocal immunofluorescent imaging of Jeg-3 WT and Myo9b KO cells. Actin signal is shown in yellow and paxillin signal in red. The boundary analysis region is shown in green and represents the cell-side band extending 600 pixels into the cell, equivalent to 30 µm. Segmented boundary-localized actin bundles are shown in blue, and segmented paxillin-positive focal adhesions are shown in purple. Scale bar 10 µm. B) Quantification of Actin Bundle Area in $\mu m^2$ from conditions in (A). Bars represent mean ± SEM; Significance by unpaired t-test (n= 47 cells for Jeg 3 WT n= 45 cells for Myo9b KO, *** =p ≤ 0.00

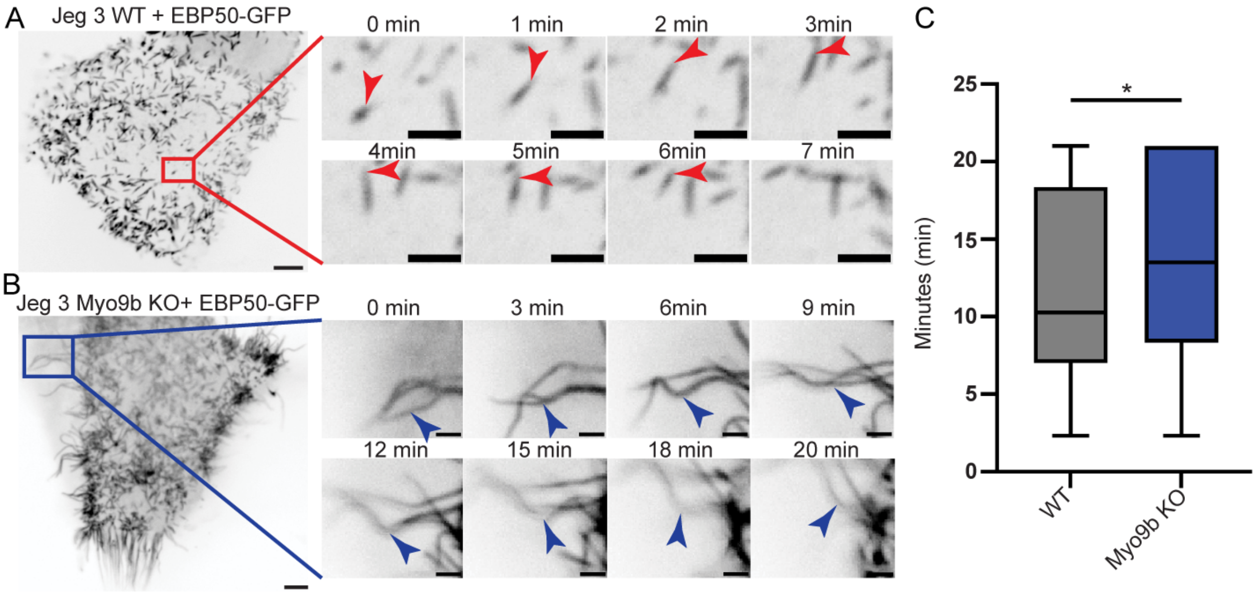


**Fig. S3**. Myo9b KO cells Contain Microvilli that Persist Longer. A) Jeg-3 WT cell transiently expressing EBP50-GFP from the first frame of Video 2, with inset images from the red box showing microvilli tracked over time (red arrow). Scale bar 5 μm; inset 2 μm B) Myo9b KO cell transiently expressing EBP50-GFP from the first frame of Video 2, with inset images from the blue box showing microvilli tracked over time (blue arrow). Scale bar 5 μm; inset 2 μm. C) Quantification of average microvilli lifetime from (A, B). No error on the upper bound of Myo9b KO cells because the movie limit for each condition was 21 minutes, and Myo9b KO cells contained many microvilli that persisted for 21 minutes. Bars represent mean ± SEM; Significance by Mann-Whitney t-test, (n=60 microvilli * =p ≤ 0.05).

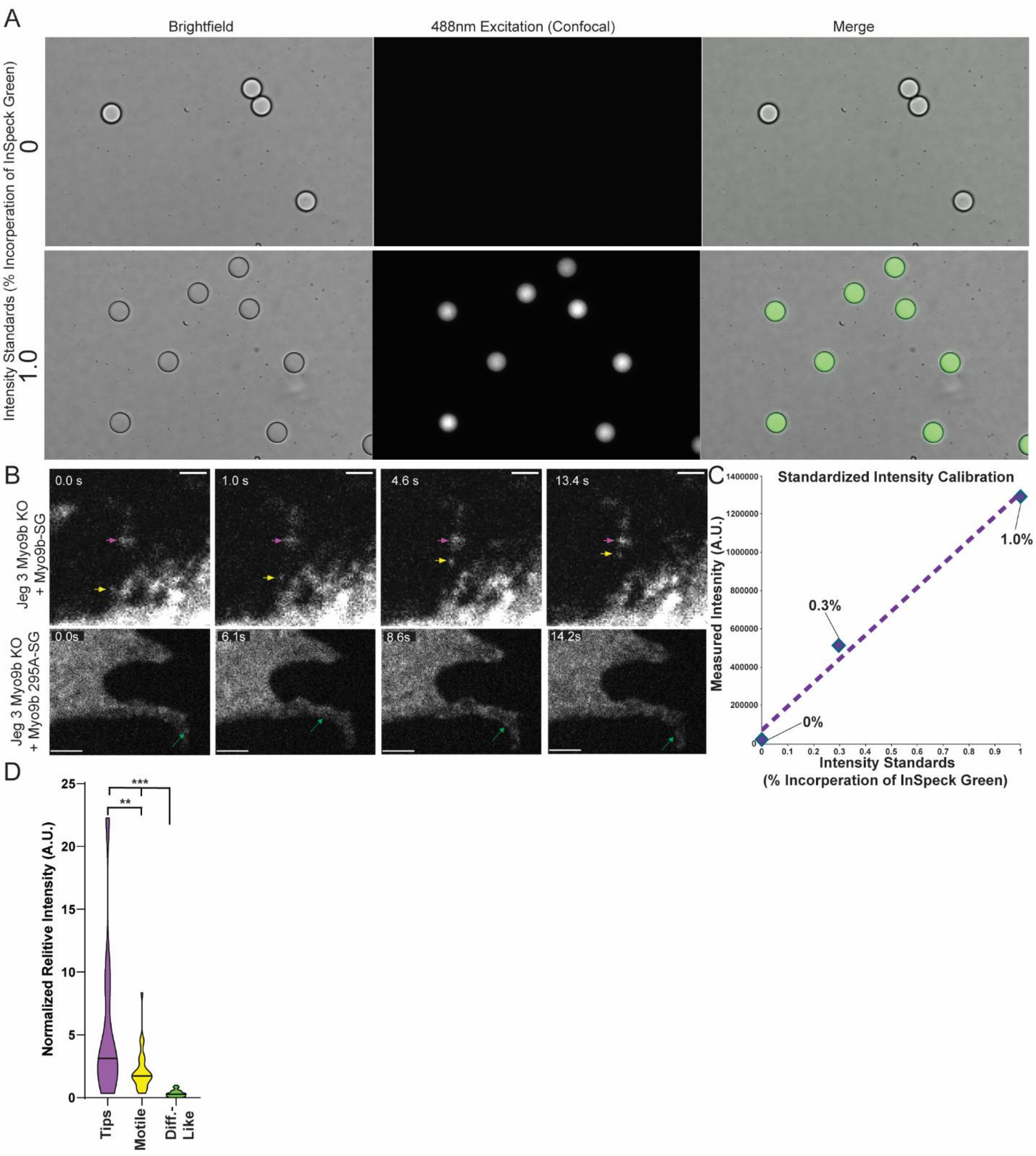


**Fig. S4**. Relative Intensity of SG-Myo9b Signal. A) InSpeck Green calibration kit 0.0% and 1.0% intensity standards imaged in brightfield and 488nm excitation (confocal). B) Image duplicated from 5A, yellow arrows indicate motile SG-Myo9b. Tip-localized SG-Myo9b is indicated by magenta arrows, and in the Myo9b KO + SG-Myo9b R295A condition, the green arrow indicates a diffusive-like particle of SG-Myo9b. Scale bar 2 μm. C) Standardized intensity calibration curve generated from the InSpeck Green calibration kit D) Normalized relative intensity of tip-localized, motile, or diffusive-like SG-Myo9b. Intensity values are relative and presented as arbitrary units (A.U.). Bars represent the distribution of intensity data, and the black line indicates the median. Significance by T-test. Tip-localized SG-Myo9b n= 11, motile SG-Myo9b n=33, diffusive like n= 23. ** =$p \leq 0.01$, *** =$p \leq 0.001$.

**Movie 1.** Myo9b Tracks With the Tips of Microvilli in Jeg-3 Cells. Spinning disk confocal live cell imaging of Myo9b KO cells transiently expressing StayGold-Myo9b (SG-Myo9b) and tdTomato-EBP50. Each frame was captured at 0.8-second intervals. Scale bar 2 μm.

**Movie 2**. Microvilli turnover in Jeg-3 WT cells. Jeg-3 WT cells transiently expressing EBP50-GFP. Each frame was taken at 30 second intervals. Scale bars 5 μm.

**Movie 3**. Myo9b KO cells contain microvilli that persist for longer. Jeg-3 Myo9b KO cells transiently expressing EBP50-GFP. Each frame was taken at 30 second intervals. Scale bars 5 μm.

**Movie 4**. Myo9b Motility in Jeg-3 Cells. Spinning disk confocal live cell imaging of Myo9b KO cells transiently expressing Stay Gold Myo9b (SG-Myo9b). The frame rate is 0.2ms. Scale bar 2 μm.

**Movie 5.** Motor ATPase activity is needed for Myo9b's motility in cells. Spinning disk confocal live cell imaging of Myo9b KO cells transiently expressing Stay Gold Myo9b R295A. Jeg-3 Myo9b KO + SG- Myo9b R295A frame rate is 0.2ms. Scale bar 2 μm.

### Myo9b Cell Lysate blots replicates

#### Rep 1

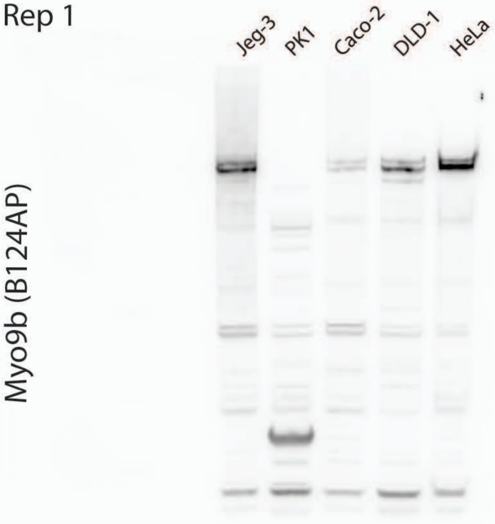


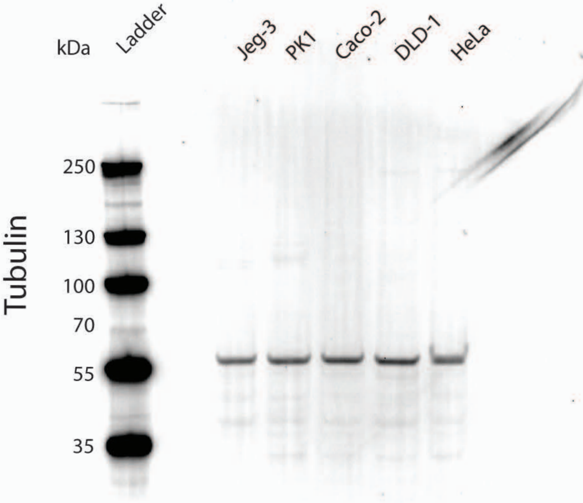


#### Rep 2

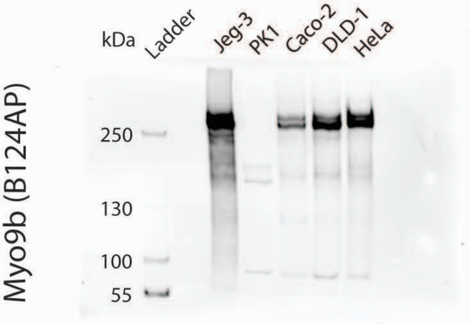


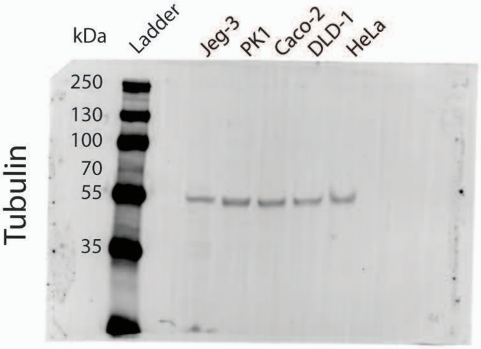


#### Rep 3

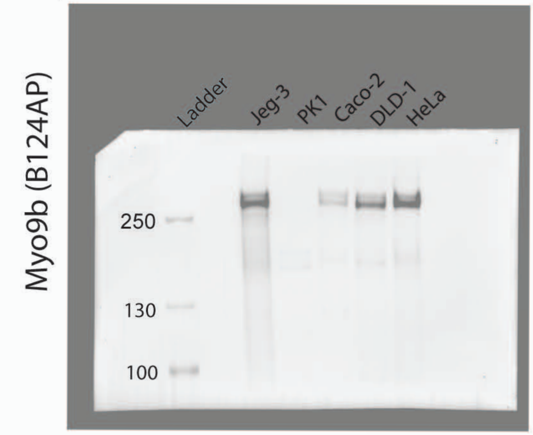


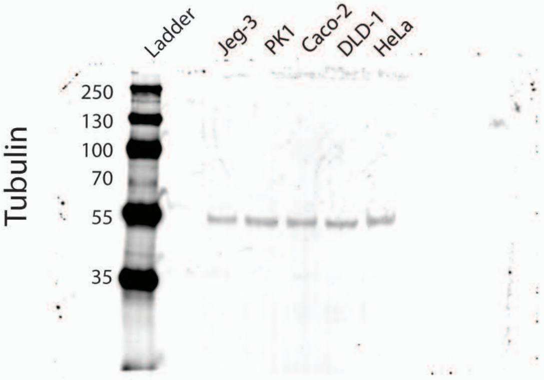

Jeg 3 Myo9b B124 AP antibody validation

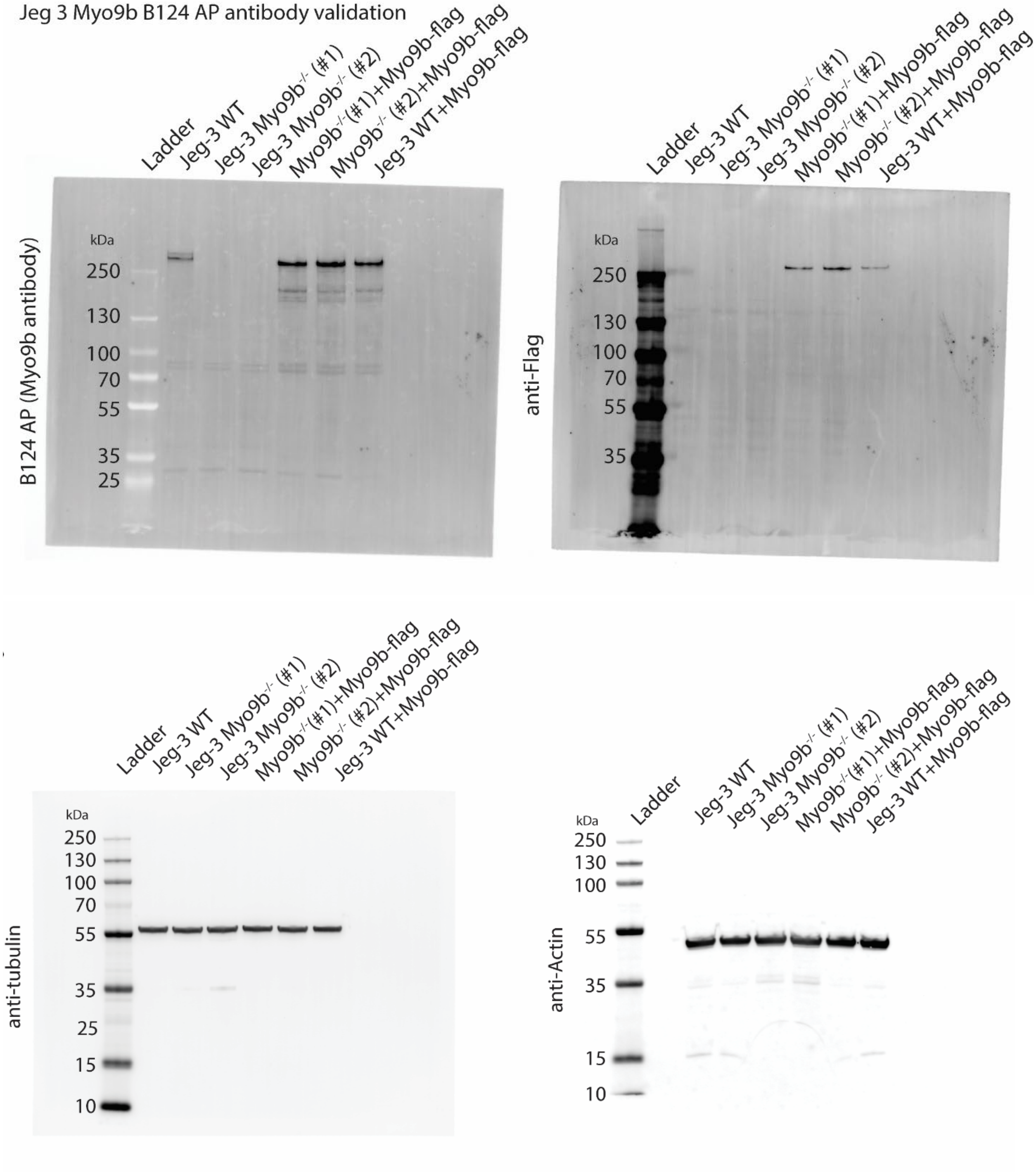
